# RNA2Immune: A Manually Curated Database of Experimentally Supported Data Linking Noncoding RNA Regulation to the Immune System

**DOI:** 10.1101/2021.12.07.471559

**Authors:** Jianjian Wang, Shuang Li, Tianfeng Wang, Si Xu, Xu Wang, Xiaotong Kong, Xiaoyu Lu, Huixue Zhang, Lifang Li, Meng Feng, Shangwei Ning, Lihua Wang

**Affiliations:** Department of Neurology, The Second Affiliated Hospital of Harbin Medical University, Harbin 150081, China; College of Bioinformatics Science and Technology, Harbin Medical University, Harbin 150081, China

**Author notes:** Corresponding authors. (Wang L), (Ning S). Equal contribution (Jianjian Wang, Shuang Li, Tianfeng Wang, Si Xu).

**Keywords:** ncRNA, Immune cells, Immune disease, Cancer immunology, Vaccine

## Abstract

Non-coding RNAs (ncRNAs), such as microRNAs (miRNAs), long noncoding RNAs (lncRNAs), circular RNAs (circRNAs), have emerged as important regulators of the immune system and are involved in the control of immune cell biology, disease pathogenesis as well as vaccine responses. A repository of ncRNA−immune associations will facilitate our understanding of ncRNA-dependent mechanisms in the immune system and advance the development of therapeutics for immune disorders as well as vaccines. Here, we describe a comprehensive database, RNA2Immune, which aims to provide a high-quality resource of experimentally supported database linking ncRNA regulatory mechanisms to immune cell function, immune disease, cancer immunology, and vaccines. The current version of RNA2Immune documents 50,433 immune−ncRNA associations in 42 host species, including: (i) 6690 ncRNA associations with immune functions involving 31 immune cell types; (ii) 38,672 ncRNA associations with 348 immune diseases; (iii) 4833 ncRNA associations with cancer immunology; and (iv) 238 ncRNA associations with vaccine responses involving 26 vaccine types targeting 22 diseases. RNA2Immune provides a user-friendly interface for browsing, searching and downloading ncRNA−immune system associations. Collectively, RNA2Immune provides important information about how ncRNAs influence immune cell function, the pathological consequences of dysregulation of these ncRNAs (immune diseases and cancers), and how ncRNAs affect immune responses to vaccines. RNA2Immune is available at http://bio-bigdata.hrbmu.edu.cn/rna2immune/home.jsp.

## Introduction

The immune system is extremely complex and versatile, simultaneously recognizing and responding to pathogens, maintaining organ homeostasis, and preventing cancer or autoimmunity [1]. Non-coding RNAs (ncRNAs), such as microRNAs (miRNAs), long non-coding RNAs (lncRNAs), circular RNAs (circRNAs) and small nucleolar RNAs (snoRNAs), have emerged as important regulators of multiple biological processes. There is now overwhelming evidence that ncRNAs play pivotal regulatory roles in the immune system [2], such as regulating immune cell development and function. For example, the *miR-212/132* cluster is enriched in hematopoietic stem cells (HSCs) and contributes to HSC differentiation and function [3], while knockout of *miR-155* affects a wide spectrum of immune responses ranging from cytokine production by T cells to antigen presentation by dendritic cells (DCs), as well as the B cell response [4]. In addition, lncRNA *Morrbid* controls the survival of neutrophils, eosinophils and monocytes in response to pro-survival cytokines by repressing *Bcl2l11* transcription [5].

Furthermore, aberrant expression of ncRNAs can contribute to a large spectrum of immune-related diseases such as autoimmune diseases, infectious diseases, hypersensitivity and graft-versus-host disease. For example, systemic lupus erythematosus (SLE) is an autoimmune disease, characterized by dysregulated innate and adaptive immune responses, which leads to chronic inflammation and tissue damage. Dysregulated expression of ncRNAs such as miRNAs (*miR-31, miR-145*), lncRNAs (*MALTAT1, ANRIL*) and circRNAs (*hsa-circ-0045272, hsa-circ-0012919*) have been associated with the pathogenesis of SLE, and are potential biomarkers and indicators of disease activity [6]. In addition, ncRNAs are involved in the development and progression of cancer by regulating immune pathways, immune cells or immune molecules. For example, *miR-200* was shown to control tumor metastasis through regulation of CD8+ tumor-infiltrating lymphocytes in non-small cell lung cancer [7]. lncRNA *LINC00240* expression was markedly increased in cervical cancer and promoted cervical cancer progression via induction of MHC class I-related chain (MIC)-mediated natural killer T cell tolerance [8].

Importantly, ncRNAs also modulate vaccine-induced host immune responses and disease prevention. Vaccines are one of the most significant inventions in improving human health, with great importance being placed on the development of biomarkers that assess the safety and efficacy of vaccine candidates. Altered miRNA expression has been linked to immunological parameters of several infectious diseases after vaccination, such as Marek’s disease [9], respiratory syncytial virus [10] and pandemic influenza [11]. Besides infectious diseases, emerging evidence indicates that miRNAs could also influence the effectiveness of cancer vaccines, such as those for breast cancer [12], colorectal cancer [13] and melanoma [14]. Moreover, miRNAs have been used in the construction of vaccines against influenza viral infection [15] and HIV infection [16]. Therefore, ncRNAs play a critical role in the development and implementation of vaccines.

To decode the role of ncRNAs in the immune system, we have established a comprehensive database, RNA2Immune, which aims to provide a high-quality resource of experimentally supported data linking ncRNAs to immune regulation across various host species. RNA2Immune provides important information about how ncRNAs influence immune cell development and function, the pathological consequences of dysregulation of these ncRNAs (such as immune diseases and cancers), and how ncRNAs modulate vaccine-mediated disease prevention. Collectively, RNA2Immune will serve as an important resource for investigating the role of ncRNAs in the immune system.

## Database framework and implementation

### Data collection and processing

To ensure a high quality database, we referred to the steps used to assemble the experimentally supported databases NSDNA [17] and Lnc2Cancer [18]. Four sections of experimentally validated datasets, including ncRNA associated with immune cell function, immune disease, cancer immunology and vaccines were collected. All data in our database was supported by experimental evidence such as RNA pull-down assay, RNAi, qRT-PCR and luciferase reporter assay. The following three steps were applied during the data collection process: i) collection of basic information, ii) searches for relevant articles on the PubMed database, and iii) extraction of relevant information from the selected articles by at least two researchers. The general workflow and features of the resource are shown in **Figure 1**.

**Figure 1.**
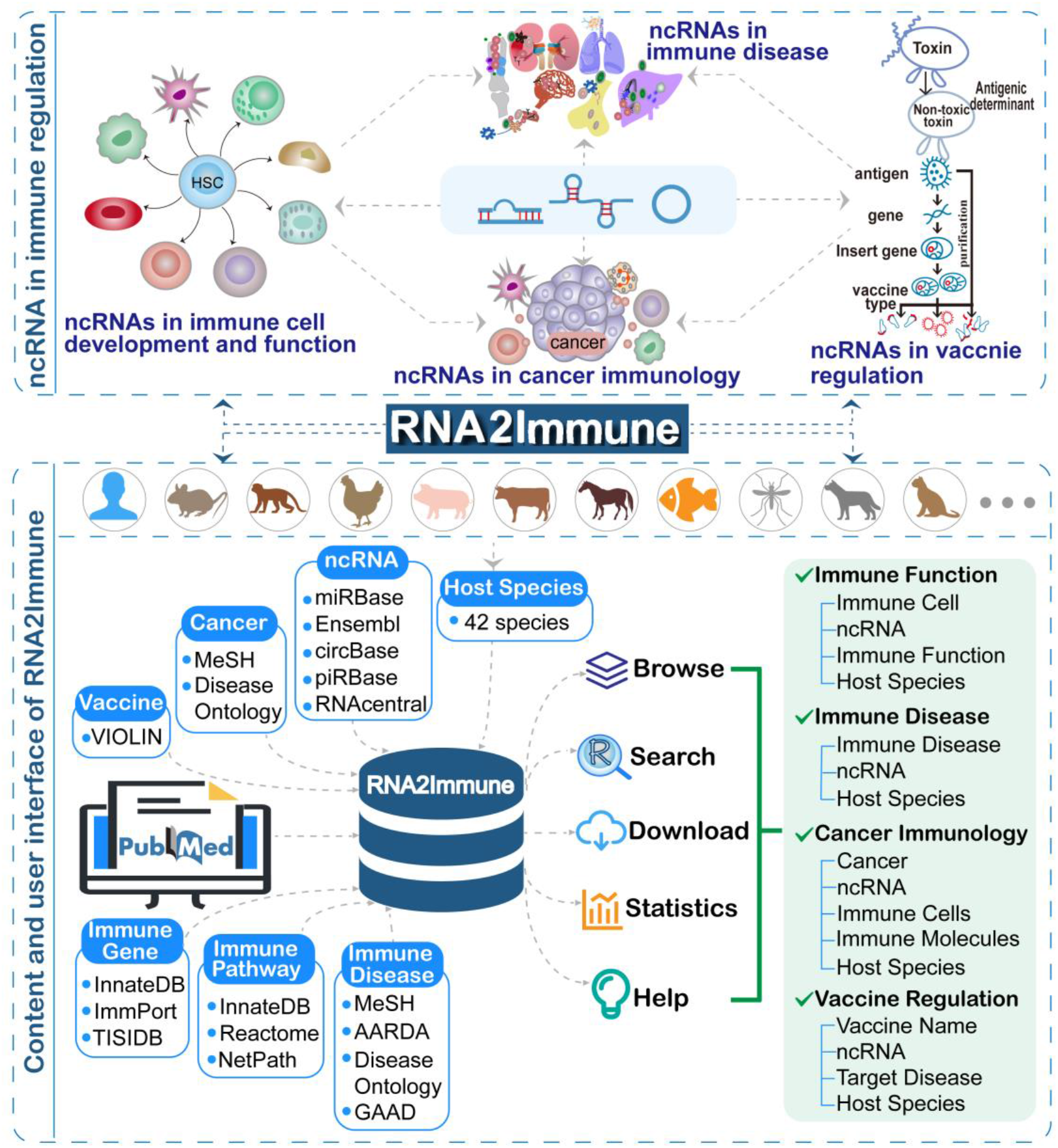
Data sources and the structure of RNA2Immune.

First, thirteen major types of ncRNA symbols were used. The ncRNA information was collected and integrated from various resources: miRNA, lncRNA, circRNA and piwi-interacting RNA (piRNA) symbols were mapped to miRBase [19], Ensembl [20], circBase [21] and piRBase [22], respectively. snoRNA, vault RNA (vtRNA), antisense RNA, small interfering RNA (siRNA), signal recognition particle (SRP) RNA, rRNA, small nuclear RNA (snRNA), Y RNA and tRNA symbols were mapped to RNAcentral [23]. The immune disease and cancer terminologies were organized based upon the controlled vocabulary of the Medical Subject Heading disease categories [24], Disease Ontology [25], American Autoimmune Related Diseases Association (AARDA, www.aarda.org) and GAAD (A Database for Autoimmune-disease Associated Genes) [26]. The immune genes were taken from InnateDB [27], ImmPort [28] and TISIDB [29]. The relevant immune pathways were extracted from the original articles and the pathway names were referenced to InnateDB [27], Reactome [30] and NetPath [31]. The annotated information about vaccines was collected from the VIOLIN (Vaccine Investigation and Online Information Network) database [32].

Second, we searched the relevant articles on the PubMed database using different keywords in each section. In the ‘Immune Function’ section, the keywords ‘each immune cell’ in combination with ‘ncRNA category name’ were used. In the ‘Immune Disease’ section, we used the keywords ‘each immune disease’ in combination with ‘ncRNA category name’. In the ‘Cancer Immunology’ section, the keywords ‘ncRNA category name’ in combination with ‘cancer,’ ‘tumor,’ ‘carcinoma,’ ‘melanoma,’ ‘sarcoma,’ ‘leukemia,’ ‘lymphoma,’ ‘myeloma,’ ‘neoplasm’ and ‘each immune gene’ or ‘each immune cell’ were used. Finally, in the ‘Vaccine’ section, we used the keywords ‘ncRNA category name’ in combination with ‘vaccine’ or ‘vaccination’.

Third, at least two researchers manually extracted relevant information from the selected articles. The general information contains the ncRNA name, genomic location and sequence of each ncRNA, immune disease (or cancer) names, host species, expression pattern of ncRNA, experimental methods, experimentally verified targets of ncRNA, detected tissue, immune pathway, a brief functional description with more detail experiment information and the corresponding literature. Moreover, each section contains specific information. The ‘Immune Function’ section, for example, contains information about the role of ncRNAs during the regulation of immune cell function events were extracted from the original articles. The categories of the ‘immune function’ were referenced to the reviews [33, 34], which include antigen presentation, cell activation, cell aging, cell apoptosis and cell self-renewal, and so on. The ‘Immune Disease’ section contains information about immune triggers and types of immunity that drive the disease. The ‘Cancer Immunology’ section contains information about ncRNA-regulated immune genes or immune cells that play critical roles in the development and progression of cancer. Finally, the ‘Vaccine’ section contains information about the vaccine name, type of vaccine, and which disease and pathogen the vaccine targets.

In addition, entries with identical ncRNA categories, ncRNA names, host species, as well as immune cell and immune functions, or disease names, or cancer names and related immune cells (molecules) or vaccine names and targeting diseases were merged into one entry, which were supported by different numbers of evidence. One example is the regulation of the macrophage inflammatory response function by *hsa-miR-21-5p*, which was reported by eight different studies. All entries from these studies were integrated into one entry.

### Database implementation

All data in the RNA2Immune database were stored and managed using the MySQL system (version 5.5). The web interface was built using Java Server Pages (JSP). The scripts for the data processing programs were written in Java. The web service runs on an Apache Tomcat web server.

### Data organization and statistics

After strict screening of more than 8700 published articles, a total of 50,433 experimentally supported associations between ncRNAs and immune associations in 42 host species were manually collected. There are four sections of experimentally supported datasets, including ncRNA associations with immune cell function, immune disease, cancer immunology and vaccines. The detailed information includes ncRNA category, ncRNA name (each ncRNA name hyperlinks to authoritative annotation databases), genomic location and sequence of each ncRNA, host species, immune disease or cancer name, expression pattern of ncRNA, experimental method, experimentally verified target of ncRNA, tissue source, immune cell, immune gene, immune function, immune pathway, immune trigger, immunity type, vaccine name, vaccine type, a brief functional description and corresponding publication information (PubMed ID, year and title of publication). The four datasets are outlined below:

#### Immune function

The ‘Immune Function’ section contains 6690 ncRNA−immune function associations from 14 host species, including 6030 miRNA, 570 lncRNA, 45 circRNA, 12 siRNA, 24 snoRNA, five piRNA, three vtRNA and one antisense RNA associations (Supplementary Table S1). This section documents 47 immune functions from 31 types of immune cells. The ten most common ncRNA−associated immune cell functions are cell differentiation, inflammatory response, cytokine production, cell proliferation, cell activation, cell apoptosis, cell polarization, cell development, antiviral response and cell migration. The five most widely studied immune cell types are macrophages, HSCs, monocytes, T cells and DCs. Among them, macrophages account for the largest number of ncRNA−immune function associations (**Figure 2**A).

**Figure 2.**
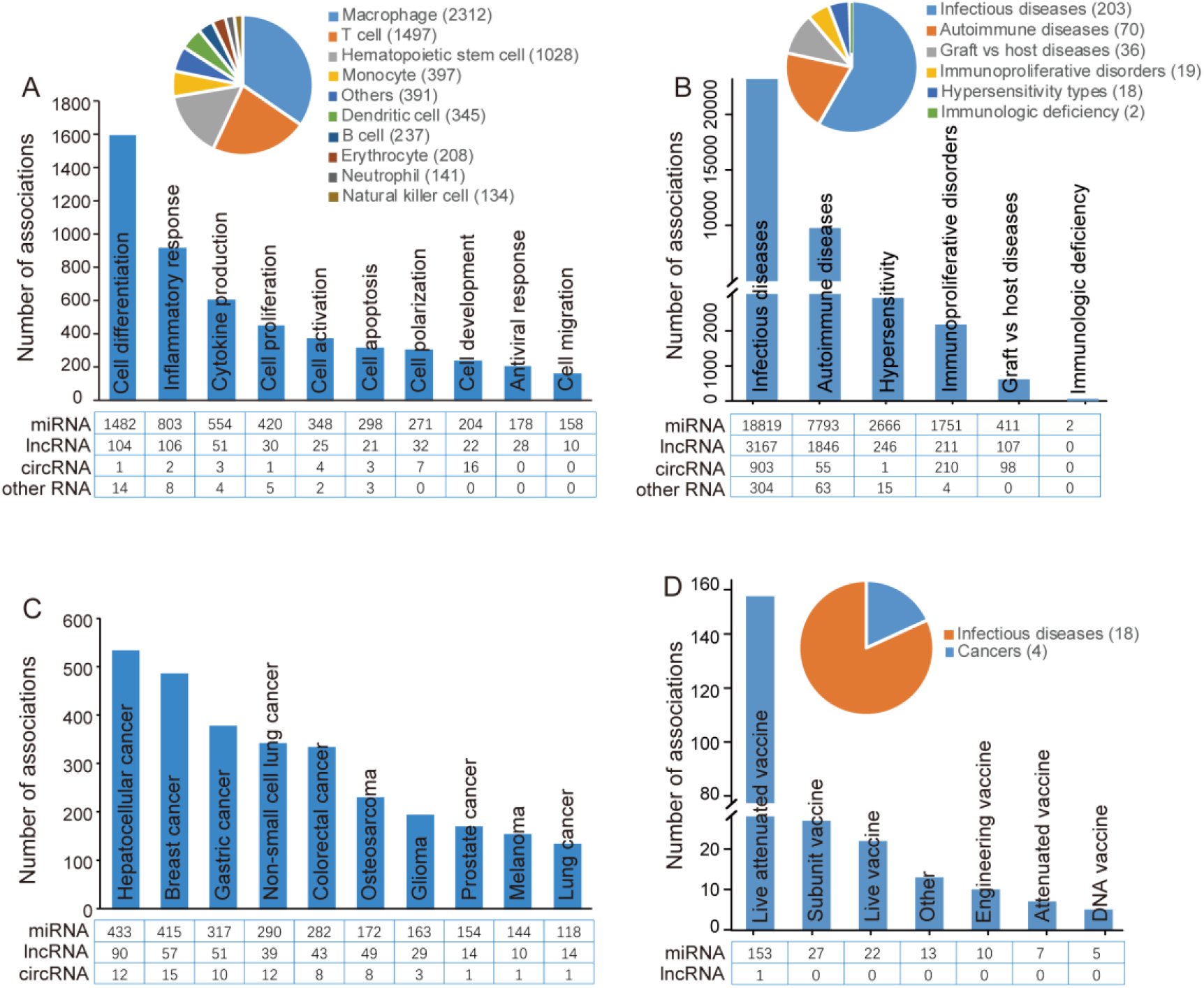
Statistics of ncRNA−immune associations in RNA2Immune. **A**. The ten most common ncRNA−immune cell functions in the ‘Immune Function’ section. The upper right pie graph shows the distribution of immune function associations for different immune cell types. **B**. The number of ncRNA−immune disease associations in different disease categories in the ‘Immune Disease’ section. The upper right pie graph shows the distribution of diseases corresponding to each category. **C**. The ten most common ncRNA−immune molecules/cells associated with cancers in the ‘Cancer Immunology’ section. **D**. The number of ncRNA−vaccine associations in different vaccine types in the ‘Vaccine’ section. The upper right pie graph shows the distribution of diseases targeting by vaccines.

#### Immune disease

The ‘Immune Disease’ section contains 38,672 ncRNA−immune disease associations across 36 host species, including 31,442 miRNA, 5577 lncRNA, 1267 circRNA, 153 siRNA, 129 snoRNA, 48 piRNA,34 tRNA, 13 snRNA, four rRNA, three SRP RNA, one vault RNA and one Y RNA associations (Supplementary Table S2). This section documents 23,193 ncRNA−infectious disease associations representing 203 infectious diseases, 9757 ncRNA−autoimmune disease associations representing 70 autoimmune diseases, 2928 ncRNA−hypersensitivity associations representing 18 hypersensitivity types, 2176 ncRNA−immunoproliferative disorder associations representing 19 immunoproliferative disorders, 616 ncRNA−graft vs host disease associations representing 36 graft vs host diseases, and two ncRNA−immunologic deficiency syndrome associations representing two immunologic deficiency syndromes (Figure 2B).

#### Cancer immunology

The ‘Cancer Immunology’ section contains 4833 ncRNA−cancer immunology associations across four host species, including 4088 miRNA, 629 lncRNA and 116 circRNA associations (Supplementary Table S3). This section documents 138 cancers, 21 immune cells and 713 immune genes, referring to 4348 immune gene associations and 485 immune cell associations. The ten most common associations between ncRNA−cancer immunology are shown in Figure 2C. Among them, hepatocellular cancer accounts for the most ncRNA−cancer immunology associations, followed by breast cancer and gastric cancer (Figure 2C).

#### Vaccine

The ‘Vaccine’ section contains 238 ncRNA−vaccine associations across nine host species, including 237 miRNA and one lncRNA associations (Supplementary Table S4). This section documents 26 vaccine types targeting 22 diseases, including 18 infectious diseases and four cancers (Figure 2D). Among them, live attenuated vaccine is the most widely studied vaccine type associated with ncRNA.

## Web interface and database utility

The RNA2Immune database is freely available at http://bio-bigdata.hrbmu.edu.cn/rna2immune/home.jsp or http://www.bio-bigdata.com/rna2immune/home.jsp. RNA2Immune provides a user-friendly web interface for an easy database query (**Figure 3**). It has been tested in Google Chrome, Mozilla Firefox and Apple Safari browsers. In addition, RNA2Immune provides a detailed tutorial in the ‘Help’ page. The main functions and application of RNA2Immune are summarized below:

**Figure 3.**
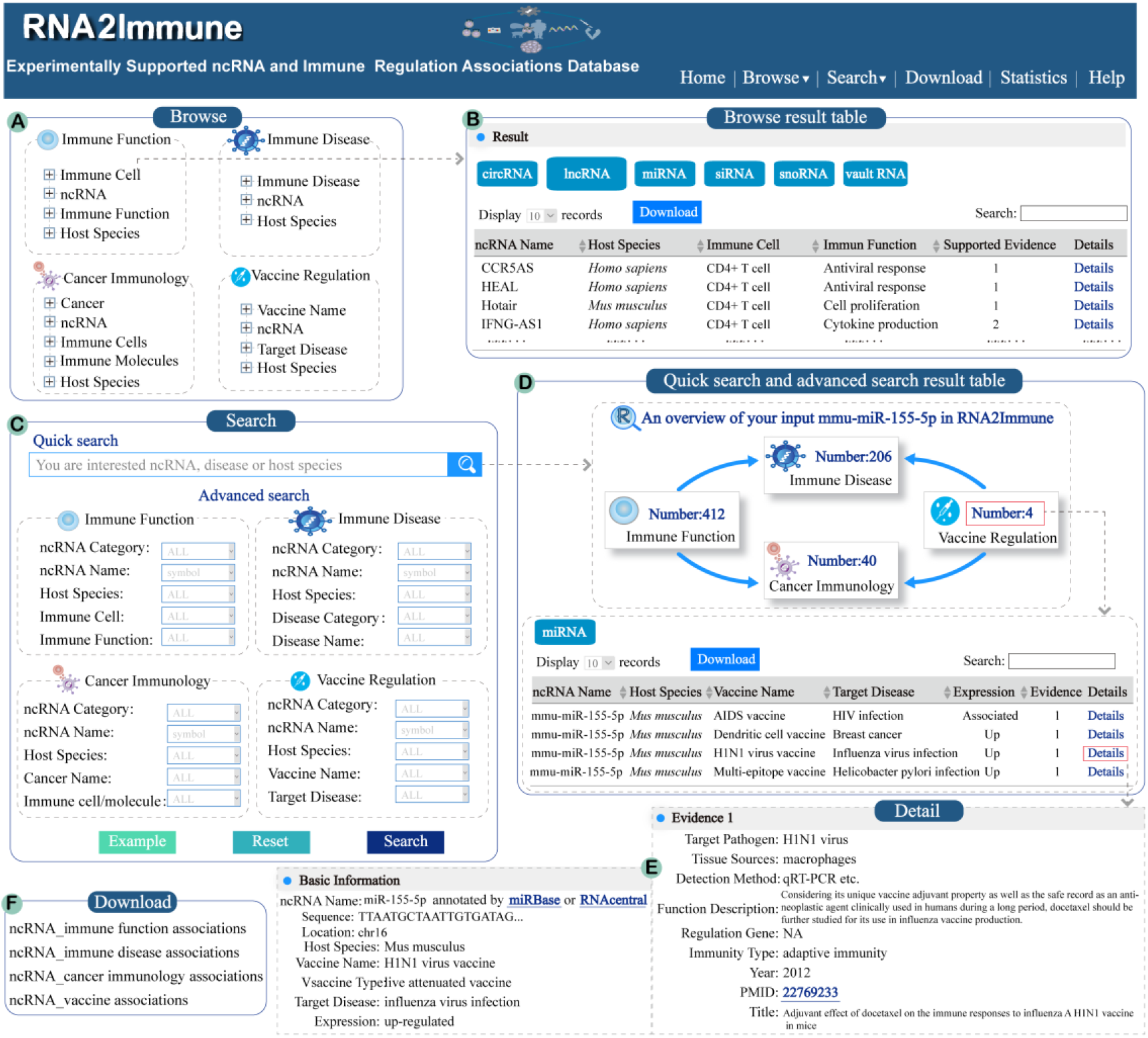
The main functions and application of RNA2Immune. **A**. Interface of the browse module. **B**. Browse result page using CD+ T cells as an example in the ‘Immune Function’ section. **C**. Interface of the ‘Quick Search’ and ‘Advanced Search’ modules. **D**. Search result page for ‘Quick Search’ and ‘Advanced Search’, using mmu-miR-155-5p as an example for the ‘Quick Search’. **E**. Interface of detailed information, using mmu-miR-155-5p in the ‘Vaccine’ section as the example. **F**. Interface of the download module.

### Browse function

The Browse page provides a catalog allowing users to browse the RNA2Immune database according to four different sections. In the ‘Immune Function’ section, users can browse by immune cells, ncRNA, immune function and host species. In the ‘Immune Disease’ section, users can browse by immune disease, ncRNA and host species. In the ‘Cancer Immunology’ section, users can browse by cancer, ncRNA, immune cells (or immune molecules) and host species. In the ‘Vaccine’ section, users can browse by vaccine name, ncRNA, targeting disease and host species. By clicking on a particular node, corresponding results are displayed in a table. Clicking the hyperlink ‘details’ allows users to obtain more detailed information about this association, including ncRNA category, ncRNA name, hyperlinks to ncRNA annotation databases, host species, immune disease or cancer name, expression pattern of ncRNA, experimental methods, experimentally verified targets of ncRNA, tissue source, immune cell, immune gene, immune function, immune pathway, immune trigger, immunity type, vaccine name, vaccine type, a brief functional description and corresponding publication information (PubMed ID, year and title of publication). In addition, RNA2Immune provides several useful features that allows the query results to be browsed easily. For example, query entries can be filtered by typing any term in the ‘Search’ box on the top right side above the table. Only entries matching the search term will be retained. Users can also click the arrows in the column headers for reordering columns in an ascending or descending manner. Users can rank the associations by the number of their supporting evidence by clicking the “Supported Evidence” button in the querying results table. Moreover, the table size per page can be selected by using the ‘Show entries’ pull-down menu on the top left side above the table. In addition, the full query result can be saved as an Excel file by clicking the ‘Download’ button on the top left side.

### Quick search

A ‘Quick Search’ interface has been developed in the home page and ‘Search’ page that allows users to search any ncRNA−immune function, ncRNA−immune disease, ncRNA−cancer immunology and ncRNA−vaccine association or all of the above. The input key words can be any ncRNA name, disease name or host species. The global search engine will present all potential results matching the key words. For example, after inputting a specific ncRNA name, users can obtain the summary of this ncRNA in the four sections of experimentally validated datasets, including immune function, immune disease, cancer immunology and vaccine. Users could further browse the entries they focused on optionally by clicking the “Number” of each section.

### Advanced search

‘Advanced Search’ provides multiple options for users to create customized searches of the four sections outlined above. Users can query the ncRNA−immune function associations of interest by combining ncRNA category, ncRNA name, host species, immune cell and immune function in the ‘Immune Function’ section. Users can query the ncRNA−immune disease associations of interest by combining ncRNA category, ncRNA name, host species, disease category and disease name in the ‘Immune Disease’ section. Users can query the ncRNA−cancer immunology associations of interest by combining ncRNA category, ncRNA name, host species, cancer name and immune cells/molecules in the ‘Cancer Immunology’ section. Users can query the ncRNA−vaccine associations of interest by combining ncRNA category, ncRNA name, host species, vaccine name and target disease in the ‘Vaccine’ section. After clicking the ‘Search’ button, the query results will be displayed in a table and can be filtered, sorted and downloaded according to the users’ needs. In addition, RNA2Immune supports fuzzy keyword searching capabilities, where all potential results matching the keywords will be presented.

### Download data

All experimentally validated ncRNA−immune cell function, ncRNA−immune disease, ncRNA−cancer immunology and ncRNA−vaccine associations can be freely accessed in the Download page. In addition, we summarized the association information for each homologous ncRNA family, which can be downloaded in the Download page.

### Examples of RNA2Immune use

As an example, the miRNA, *mmu-miR-155-5p*, was used as an input in the ‘Quick Search’ interface. Based on this input, users would obtain an overview of *mmu-miR-155-5p* in RNA2Immune, including 412 immune cell function associations, 206 immune disease associations, 40 cancer immunology associations and four vaccine associations (**Figure 4**A). After carefully reviewing these associations, we observed some important regulatory clues. For example, entries can be filtered by typing ‘antiviral response’ in the ‘Search’ box in the immune cell function associations. By doing this, we found that *miR-155-5p* is involved in the regulation of the antiviral response of T cells (CD8+ T cells and memory T cells), macrophages and natural killer cells (Figure 4B). RNA2Immune identified studies in which *miR-155-5p* was shown to contribute to the pathogenesis of viral infectious diseases like influenza virus infection [35] (Figure 4C) and *miR-155-5p* affects the immune response to the influenza A H1N1 vaccine in mice [36]. In addition, evidence for a critical role for miR-155 in driving an antitumor response in breast cancer via regulation of DCs was found by RNA2Immune, as well as reports that overexpression of *miR-155* enhances the efficacy of a DC vaccine against breast cancer in mice [37, 38] (Figure 4D–F). Another example in ‘Immune Disease’ section in advanced search page, by using the keyword combinations of ‘*hsa-miR-146a-5p*’ and ‘*Homo sapiens*’ and ‘hepatitis B’, all corresponding results are displayed. By clicking the “details”, users could further obtain detailed information about this association. Thus, RNA2Immune provides a global perspective on ncRNA functions in the immune system, ranging from immune cell biology and disease pathogenesis to vaccine responses. We look forward to seeing more users confirm and consolidate the usefulness of this database.

**Figure 4.**
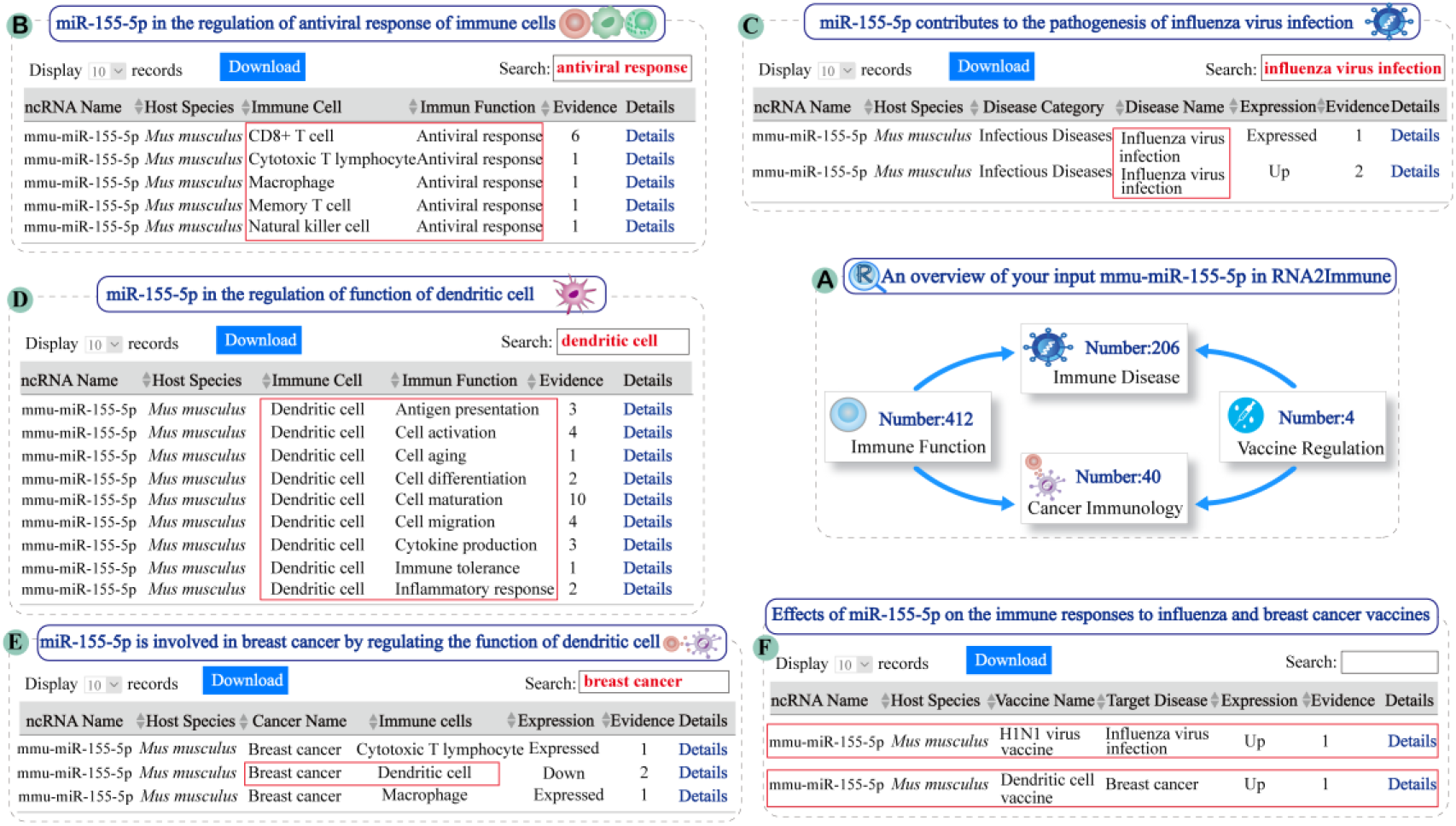
An example of RNA2Immune use. **A**. An overview of the input ‘mmu-miR-155-5p’ into the RNA2Immune database. **B**. miR-155-5p in the regulation of the antiviral response of immune cells. **C**. miR-155-5p contributes to the pathogenesis of the influenza virus infection. **D**. miR-155-5p is involved in the regulation of DC function. **E**. miR-155-5p is involved in breast cancer by regulating the function of DCs. **F**. Effects of miR-155-5p on the immune responses to the influenza and breast cancer vaccines.

## Discussion and perspectives

Recently, great effort has been made to develop beneficial databases for exploring the complex immune system. For example, IEDB (Immune Epitope Database) [39] focuses on the immune epitope data, while the DICE project (Database of Immune Cell Expression, Expression quantitative trait loci (eQTLs) and Epigenomics) [40] provides polymorphisms of immune-related genes. Immuno-Navigator (a batch-corrected gene expression and coexpression database for immune cell types) [41] contain information about gene co-expression in immune cells. VIOLIN (Vaccine Investigation and Online Information Network), a web-based vaccine database, integrates vaccine literature data mining, vaccine research data curation and storage, and curated vaccine data analysis [32]. These databases are extremely valuable to advance our mechanistic understanding of immunology. However, to the best of our knowledge, none of these resources were developed to specifically collect data on the role of ncRNAs in the immune system across various species.

ncRNA regulation has emerged as a critical regulatory component in immune system development, immune response and disease prevention [4]. With the development of new approaches and technologies, there has been an explosion of data about regulatory ncRNAs in the immune system in the past decade, ranging from immune cell biology and disease pathogenesis to vaccine response. However, these data are dispersed in thousands of independent studies. Therefore, there is an urgent need to collect experimentally supported immune−ncRNA associations to advance our understanding of immunology, and facilitate the development of vaccines and therapeutics for disorders like infection, autoimmune diseases, and cancer. Thus, to meet this need, we have established a comprehensive database, RNA2Immune, which houses a manually curated collection of experimentally supported immune−ncRNA associations across 42 host species.

The current version of RNA2Immune documents 50,433 experimentally validated ncRNA−immune associations, including: (i) 6690 ncRNA−immune function associations from 14 host species; (ii) 38,672 ncRNA−immune disease associations across 36 host species; (iii) 4483 ncRNA−cancer immunology associations, referring to 4348 immune molecule associations and 485 immune cells associations in 138 cancers; (iv) 238 ncRNA−vaccine associations in nine host species. For any specific ncRNA, RNA2Immune can provide global evidence for this ncRNA in the regulation of immune cell function, pathological consequences of dysregulation of this ncRNA (such as immune diseases and cancers), and immune responses to vaccines against diseases.

In conclusion, the role of ncRNAs are emerging as critical regulators of the immune system. RNA2Immune focuses on ncRNA-mediated regulation of the immune system in the context of immune cell function, immune disorder, cancer and vaccination. With the increasing interest in ncRNA-mediated regulation of biological processes, we hope that RNA2Immune will serve as a valuable resource for helping researchers to facilitate the understanding of ncRNA in the regulation of the immune system. The development of high-throughput technologies such as RNA-seq or microarray, has produced more extensive data linking ncRNA and immune system. These advances will offer the opportunity to extend the RNA2Immune. In the future, we will continue to update the database and website in regular intervals per year depending on the increasing numbers of new studies focusing on immune-related ncRNAs. We also plan to integrate high-throughput data and more functional annotations resources, and provide additional tools such as a disease association prediction tool.

## Data availability

The RNA2Immune database is freely available at http://bio-bigdata.hrbmu.edu.cn/rna2immune/home.jsp or http://www.bio-bigdata.com/rna2immune/home.jsp.

## CRediT author statement

**Jianjian Wang:** Software, Data curation, Methodology, Writing - Original Draft. **Shuang Li:** Data curation, Methodology, Software, Formal analysis. **Tianfeng Wang:** Data curation, Methodology, Software, Formal analysis. **Si Xu:** Data curation, Methodology, Software, Formal analysis. **Xu Wang:** Data curation. **Xiaotong Kong:** Data curation. **Xiaoyu Lu:** Data curation. **Huixue Zhang:** Data curation. **Lifang Li:** Formal analysis. **Meng Feng:** Formal analysis. **Shangwei Ning:** Conceptualization, Project administration, Writing - review & editing. **Lihua Wang:** Conceptualization, Project administration, Writing - review & editing.

## Acknowledgments

This work was supported by the National Natural Science Foundation of China (Grant Nos. 81820108014, 82071407, 81771361, 32070672 and 81901277), National Key R&D Project (Grant No. 2018YFE0114400), The Postdoctoral Foundation of Heilongjiang Province (Grant No. LBH-TZ1019).

## Supplementary material

**Table S1 Statistics for the immune function−ncRNA associations in different host species in the RNA2Immune database**

**Table S2 Statistics for the immune disease−ncRNA associations in different host species in the RNA2Immune database**

**Table S3 Statistics for the cancer immunology−ncRNA associations in different host species in the RNA2Immune database**

**Table S4 Statistics for the vaccine−ncRNA associations in different host species in the RNA2Immune database**

