## Supplementary material for "RNA2Immune: A Manually Curated Database of Experimentally Supported Data Linking Noncoding RNA Regulation to the Immune System": Table S1

Table S1 Statistics for the immune function−ncRNA associations in different host species in the RNA2Immune database

| Species | miRNA | lncRNA | circRNA | siRNA | snoRNA | piRNA | vault RNA | atRNA | Total |
| --- | --- | --- | --- | --- | --- | --- | --- | --- | --- |
| Homo sapiens | 2742 | 269 | 25 | 6 | 20 | 5 | 3 | - | 3070 |
| Mus musculus | 3087 | 272 | 20 | 6 | 4 | - | - | 1 | 3390 |
| Danio rerio | 27 | - | - | - | - | - | - | - | 27 |
| Drosophila melanogaster | 4 | - | - | - | - | - | - | - | 4 |
| Gallus gallus | 22 | 6 | - | - | - | - | - | - | 28 |
| Rattus norvegicus | 69 | 3 | - | - | - | - | - | - | 72 |
| Sus scrofa | 30 | 12 | - | - | - | - | - | - | 42 |
| Bos taurus | 9 | - | - | - | - | - | - | - | 9 |
| Miichthys miiuy | 11 | - | - | - | - | - | - | - | 11 |
| Gadus morhua | 8 | - | - | - | - | - | - | - | 8 |
| Macaca nemestrina | 19 | - | - | - | - | - | - | - | 19 |
| Marsupenaeus japonicus | 1 | - | - | - | - | - | - | - | 1 |
| Canis lupus familiaris | 1 | - | - | - | - | - | - | - | 1 |
| Cyprinus carpio | - | 8 | - | - | - | - | - | - | 8 |
| Total | 6030 | 570 | 45 | 12 | 24 | 5 | 3 | 1 | 6690 |
