## Supplementary material for "RNA2Immune: A Manually Curated Database of Experimentally Supported Data Linking Noncoding RNA Regulation to the Immune System": Table S2

Table S2 Statistics for the immune disease−ncRNA associations in different host species in the RNA2Immune database

| Species | miRNA | lncRNA | circRNA | siRNA | snRNA | snoRNA | piRNA | rRNA | tRNA | Y RNA | vault RNA | SRP RNA | Total |
| --- | --- | --- | --- | --- | --- | --- | --- | --- | --- | --- | --- | --- | --- |
| Homo sapiens | 20315 | 4127 | 1029 | 73 | 13 | 123 | 19 | 1 | 21 | 1 | 1 | 3 | 25726 |
| Mus musculus | 5883 | 1021 | - | 50 | - | 3 | - | - | - | - | - | - | 6957 |
| Sus scrofa | 1064 | 110 | 131 | 6 | - | 2 | - | 1 | 10 | - | - | - | 1324 |
| Gallus gallus | 1068 | 102 | 87 | 4 | - | - | - | - | - | - | - | - | 1261 |
| Rattus norvegicus | 688 | 144 | - | 7 | - | 1 | - | - | - | - | - | - | 840 |
| Bos taurus | 671 | - | - | - | - | - | - | - | 2 | - | - | - | 673 |
| Macaca mulatta | 269 | - | - | - | - | - | - | 2 | - | - | - | - | 271 |
| Aedes aegypti | 186 | 6 | - | 6 | - | - | - | - | 1 | - | - | - | 199 |
| Aedes albopictus | 134 | - | - | - | - | - | 25 | - | - | - | - | - | 159 |
| Anopheles sinensis | 148 | - | - | - | - | - | - | - | - | - | - | - | 148 |
| Felis catus | 143 | - | - | 5 | - | - | - | - | - | - | - | - | 148 |
| Cyprinus carpio | 120 | - | - | - | - | - | - | - | - | - | - | - | 120 |
| other | 753 | 67 | 20 | 2 | - | - | 4 | - | - | - | - | - | 846 |
| Total | 31442 | 5577 | 1267 | 153 | 13 | 129 | 48 | 4 | 34 | 1 | 1 | 3 | 38672 |
