## Supplementary material for "RNA2Immune: A Manually Curated Database of Experimentally Supported Data Linking Noncoding RNA Regulation to the Immune System": Table S3

Table S3 Statistics for the cancer immunology−ncRNA associations in different host species in the RNA2Immune database

| Species | miRNA | lncRNA | circRNA | Total |
| --- | --- | --- | --- | --- |
| Homo sapiens | 3697 | 583 | 105 | 4385 |
| Mus musculus | 369 | 45 | 11 | 425 |
| Rattus norvegicus | 20 | 1 | - | 21 |
| Canis lupus familiaris | 2 | - | - | 2 |
| Total | 4088 | 629 | 116 | 4833 |
