## Supplementary material for "RNA2Immune: A Manually Curated Database of Experimentally Supported Data Linking Noncoding RNA Regulation to the Immune System": Table S4

Table S4 Statistics for the vaccine−ncRNA associations in different host species in the RNA2Immune database

| Species | miRNA | lncRNA | Total |
| --- | --- | --- | --- |
| Homo sapiens | 113 | - | 113 |
| Mus musculus | 72 | 1 | 73 |
| Gallus gallus | 21 | - | 21 |
| Sus scrofa | 21 | - | 21 |
| Ovis aries | 3 | - | 3 |
| Bos taurus | 2 | - | 2 |
| Macaca mulatta | 2 | - | 2 |
| Oncorhynchus mykiss | 2 | - | 2 |
| Paralichthys olivaceus | 1 | - | 1 |
| Total | 237 | 1 | 238 |
